# Bcr-Abl tyrosine kinase inhibitor imatinib as a potential drug for COVID-19

**DOI:** 10.1101/2020.06.18.158196

**Authors:** Nirmitee Mulgaonkar, Haoqi Wang, Samavath Mallawarachchi, Sandun Fernando, Byron Martina, Daniel Ruzek

## Abstract

The rapid geographic expansion of severe acute respiratory syndrome coronavirus 2 (SARS-CoV-2), the infectious agent of Coronavirus Disease 2019 (COVID-19) pandemic, poses an immediate need for potent drugs. Enveloped viruses infect the host cell by cellular membrane fusion, a crucial mechanism required for virus replication. The SARS-CoV-2 spike glycoprotein, due to its primary interaction with the human angiotensin-converting enzyme 2 (ACE2) cell-surface receptor, is considered as a potential target for drug development. Based on *in silico* screening followed by *in vitro* studies, here we report that the existing FDA-approved Bcr-Abl tyrosine kinase inhibitor, imatinib, inhibits SARS-CoV-2 with an IC_50_ of 130 nM. We provide evidence that although imatinib binds to the receptor-binding domain (RBD) of SARS-CoV-2 spike protein with an affinity at micromolar, i.e., 2.32 ± 0.9 μM levels, imatinib does not directly inhibit the spike RBD:ACE2 interaction – suggesting a Bcr-Abl kinase-mediated fusion inhibition mechanism is responsible for the inhibitory action. We also show that imatinib inhibits other coronaviruses, SARS-CoV, and MERS-CoV via fusion inhibition. Based on promising *in vitro* results, we propose the Abl tyrosine kinase inhibitor (ATKI), imatinib, to be a viable repurposable drug against COVID-19.

## Introduction

In early December 2019, the Chinese health authorities reported several cases of pneumonia of unknown cause that had originated in Wuhan, a city in the Hubei province of China. The causative agent of this outbreak was identified to be a virus that belonged to the *Sarbecovirus* subgenus, *Orthocoronavirinae* subfamily which was previously referred to by its interim name 2019 novel coronavirus (2019-nCoV) [1, 2] and was later named as SARS-CoV-2 [3]. Due to the rapid spread of COVID-19, the World Health Organization (WHO) declared it a global pandemic in March 2020 [4]. By mid-August 2020, over 21 million cases have been confirmed around the world, resulting in more than 768,000 deaths [5]. Unfortunately, there is no approved antiviral treatment or preventive vaccine for coronaviruses in humans. Since supportive care is the only recommended interim treatment, it is imperative to identify repurposable lead compounds to rapidly treat COVID-19 patients until a SARS-CoV-2-specific drug and a vaccine is developed.

Although the coronavirus genome consists of numerous conserved druggable enzymes, including papain-like protease (PLpro), 3C-like protease (3CLpro), non-structural proteins RNA-dependent RNA polymerase (RdRp) and helicase, development of clinically approved antiviral therapies has proven to be a difficult task [6]. The surface structural spike glycoprotein (S), a key immunogenic CoV antigen essential for virus and host cell-receptor interactions, is an important target for therapeutic development. The spike protein consists of an N-terminal S1 subunit (receptor binding) and a C-terminal S2 subunit (membrane fusion). The S1 subunit contains the receptor-binding domain (RBD) which attaches to the host membrane, thus playing an important role in viral entry. SARS-CoV-2 utilizes the ACE2 receptor for entry and the transmembrane protease, serine 2 (TMPRSS2) for spike protein priming [7]. Crystallographic studies have shown that SARS-CoV-2 binds to the ACE2 receptor, with a binding mode nearly identical to that of SARS-CoV [8-11]. The binding affinity of the ACE2 receptor to the RBD of the SARS-CoV-2 spike protein is reported to be significantly higher as compared to SARS-CoV [10, 11].

Based on the importance of virus membrane fusion events in the viral life cycle and its infectivity, the spike protein of SARS-CoV-2 was targeted for drug screening. This study utilizes *in silico* methodology followed by in vitro experimental validation to screen existing FDA-approved small molecule drugs specific to the RBD of the spike protein of SARS-CoV-2 to identify repurposable drugs targeting further clinical validation.

## Results

### Protein Structure Prediction and Validation

A model for SARS-CoV-2 spike protein was constructed using the crystal structure (6VSB_chain A) to correct missing residues. The amino acid sequence identity between the target sequence (GenBank: QHD43416.1) and template (6VSB_chain A) was 99.58%. The SARS-CoV-2 model showed an RMSD of 0.4683 Å relative to the crystal structure (6VSB_chain A). Structure assessment of the predicted model using the Ramachandran plot showed 90.04% residues in the most favored regions with 2.03% outliers. None of the outliers contained the residues present at the active site of the protein. The predicted model was further used for *in-silico* studies.

### Ligand Screening

In virtual screening, a library of approximately 5,800 compounds was docked against the SARS-CoV-2 spike RBD protein. The output was analyzed for common classes of drugs with highest (most negative) docking scores that resulted in seven compounds with three compounds, Antiviral825, Antiviral2038 and Antiviral2981 with docking scores of −6.30 ± 0.00, −6.20 ± 0.20 and −6.00 ± 0.00 kcal/mol from the Enamine Antiviral library, and four compounds, ponatinib, imatinib, ergotamine, and glecaprevir with docking scores of −7.63 ± 0.06, −6.80 ± 0.17, −7.70 ± 0.00 and −7.20 ± 0.35 kcal/mol from the ZINC15 FDA library respectively. The above libraries were chosen to help identify a repurposable drug that can potentially inhibit the SARS-CoV-2. The screened compounds had the highest scores within their respective sets and had one or more binding conformations at the ACE2 binding domain of the spike protein. The most common class of drugs was found to be Abl tyrosine kinase inhibitors (ATKI), and hence two drugs (ponatinib and imatinib) with the highest scores were selected for *in vitro* testing. The binding scores for the seven screened compounds at the RBD are shown in Fig. 1A and detailed description of the screened drugs is given in Table S2 under Supplementary Data. The high affinity of the screened compounds is visible when compared with the negative control dimethyl sulfoxide (DMSO), which is ineffective against coronaviruses [12].

**Fig. 1.**
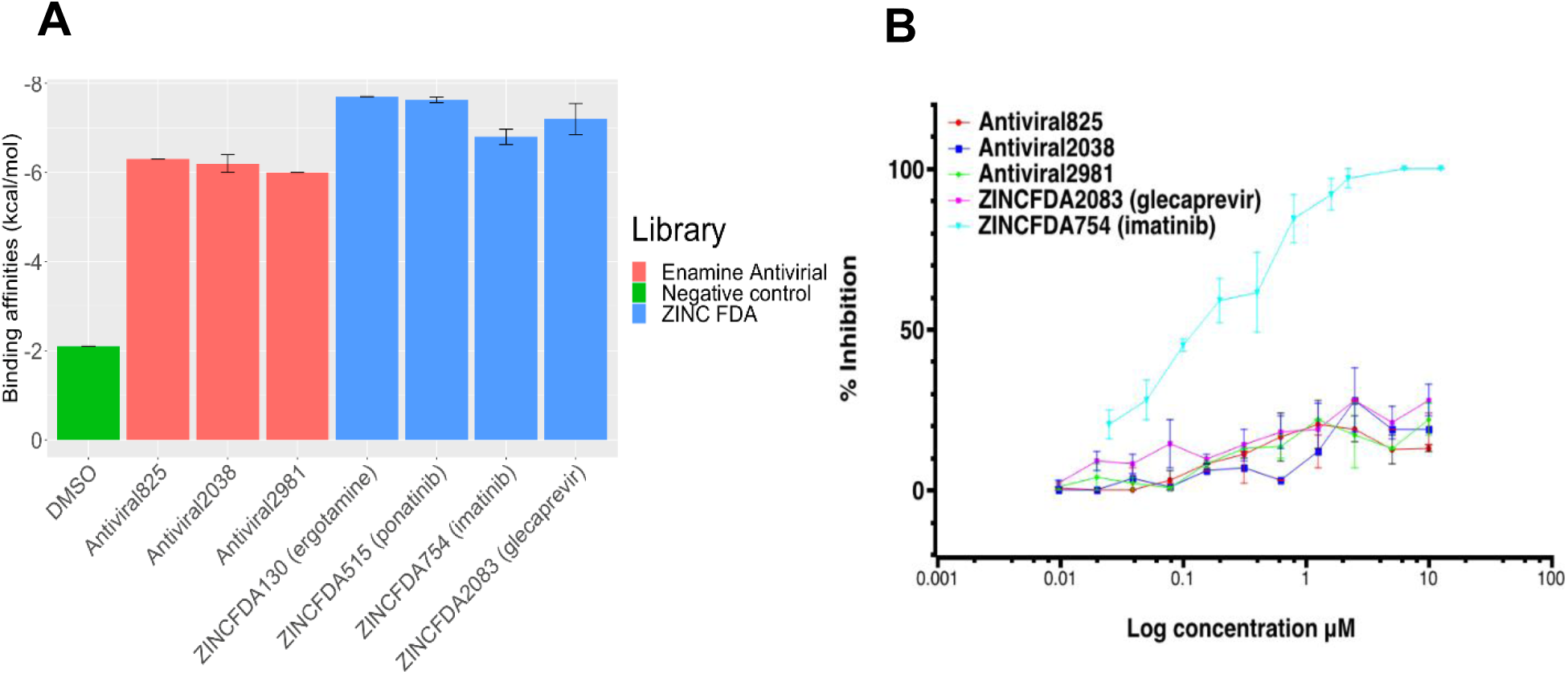
**A]** Binding scores of the selected compounds and DMSO (negative control) and, **B]** Percent inhibition compared to the amount of plaques on cells. Here, 2-fold dilution of the compounds were done in duplo. Then 300 TCID50 of SARS-CoV-2 was added to each well, and plates were incubated at 37C for 1 hour. Then, the mixes were added onto vero cells and incubated for 8 hours at 37 C. Subsequently, cells were fixed for 15 min with 2% PFA, followed by another 15 min fixation with 70% ethanol. Fixed cells were stained with a monoclonal antibody, followed by AlexaFluor 488. Here imatinib shows significant % inhibition as compared to other compounds tested.

Based on promising *in silico* data, and initial viral plaque assay results (Fig. 1A), imatinib was chosen to be advanced for further experimental validation. (Due to a supply-shortage ponatinib and ergotamine were unavailable for purchase and hence could not be included in the initial viral plaque assays). Previous studies have shown imatinib to inhibit SARS-CoV and MERS-CoV by blocking endosomal fusion at the cell-culture level [6, 12-15]. It has been suggested that tyrosine-kinase inhibitors do not affect the cleavage of the spike protein but inhibit spike-mediated endosomal fusion [6, 12, 13]. The high affinity of tyrosine-kinase inhibitors towards the spike protein is deduced from the initial docking results, where both imatinib and ponatinib have shown highly negative binding free energies.

### *In vitro* efficacy of imatinib

First, we evaluated the toxicity of imatinib when incubating the compound on Vero cells for one hour or eight hours. In the experiments where the compound remained on the cells for eight hours toxicity was measured at concentrations of 25 μM, 12.5 μM, 6.3 μM, and 3.2 μM. However, in the 1-hour design, no toxicity was observed. Next, we evaluated the ability of imatinib to inhibit replication and entry. At concentrations as low as 0.2 μM the compound was effective in suppressing 50% of plaque formation in the 8-hr design, and the IC_50_ value determined using linear regression was 130 nM. Consistent with the toxicity data, toxicity was observed between 25 and 3.2 μM. The compound also showed efficacy in the 1-hour design, with higher IC_50_ values. These data indicate that imatinib inhibits virus replication *in vitro* as shown in Fig. 2A and 2B.

**Fig. 2.**
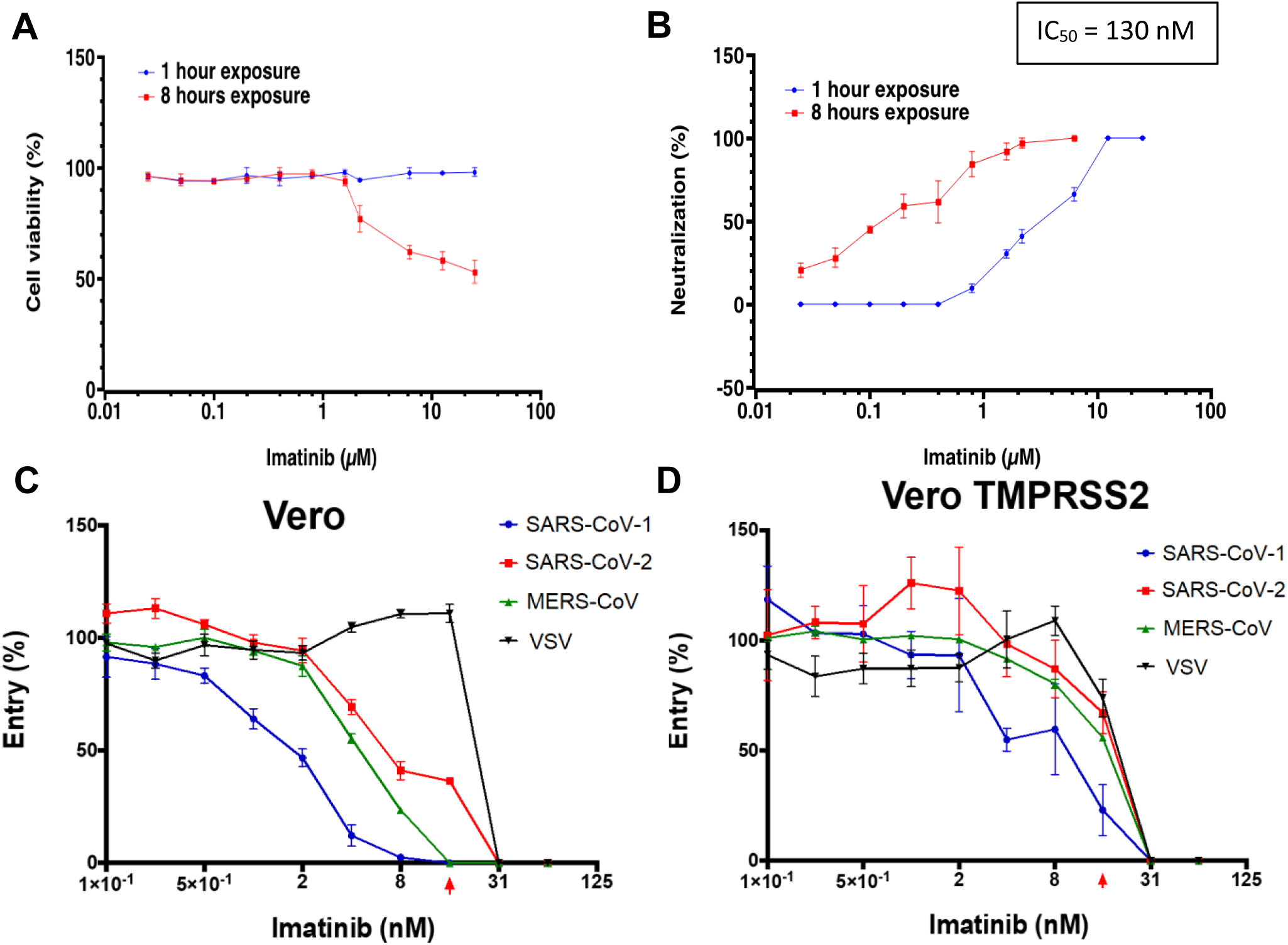
**A]** Cell viability after incubation of Vero cells with imatinib for either 1 or 8 hours. and **B]** SARS-CoV-2 neutralization profile post 1- and 8-hours exposure to imatinib. Inhibition of VSV pseudoparticles for SARS-CoV, SARS-CoV-2, MERS-CoV and VSV(control) after incubation with imatinib in **C]** Vero cells and **D]** Vero-TMPRSS2 cells. The red arrow indicates the concentration where no toxicity was observed microscopically anymore (15 nM).

### Imatinib inhibits fusion

To evaluate if imatinib inhibits viral entry, we performed two fusion assays: endosomal (Vero) and plasma membrane (Vero-TMPRSS2) as shown in Fig. 2C and 2D, respectively. Based on cytotoxicity, at concentrations below 15 nM, no toxicity was observed microscopically (red arrow in the graph). VSV-G control revealed 100% infectivity (cytopathic effect at every concentration below this, suggesting the inhibitor did not affect VSV-G entry. VSV-G particles cells do not carry spike proteins and thus, no significant entry inhibition occurred, suggesting that entry inhibition is likely mediated through the spike protein. However, the effect on Vero-TMPRSS2 cells was less clear for any of the coronaviruses used when compared to the VSV-G control. A similar level of toxicity was observed in these cells. It is worth noting that toxicity is probably the result of incubating cells with imatinib for 16 hours in the assay. Taken together, there is evidence that imatinib inhibits spike fusion and prevents viral entry, possibly by preventing endosomal entry.

### Binding kinetics analyses via Biolayer Interferometry (BLI)

The binding kinetics of imatinib to the RBD of SARS-CoV-2 spike protein was evaluated using biolayer interferometry (BLI), as shown in Fig. 3A. The analysis showed that imatinib binds to the SARS-CoV-2 RBD protein with an on-rate (k_on_) as (3.22 ± 0.45) × 10^3^ M^-1^ s^-1^ and dissociates with an off-rate (k_off_) as (7.07 ± 1.87) × 10^−3^ s^-1^. This resulted in an equilibrium affinity constant (K_D_) 2.32 ± 0.9 μM which is calculated as a ratio of the k_off_ and k_on_ rates. The affinity values indicate that 50% of the RBDs on the surface spike glycoproteins will be occupied at micromolar concentrations of imatinib. However, this value is too close to the toxicity levels observed in the above assays and very high compared to the IC_50_ value, as well as the nanomolar affinity of ACE2 on immobilized RBD (Fig. S1) suggest that it is likely to inhibit spike fusion by the other previously suggested MOA [12].

**Fig. 3.**
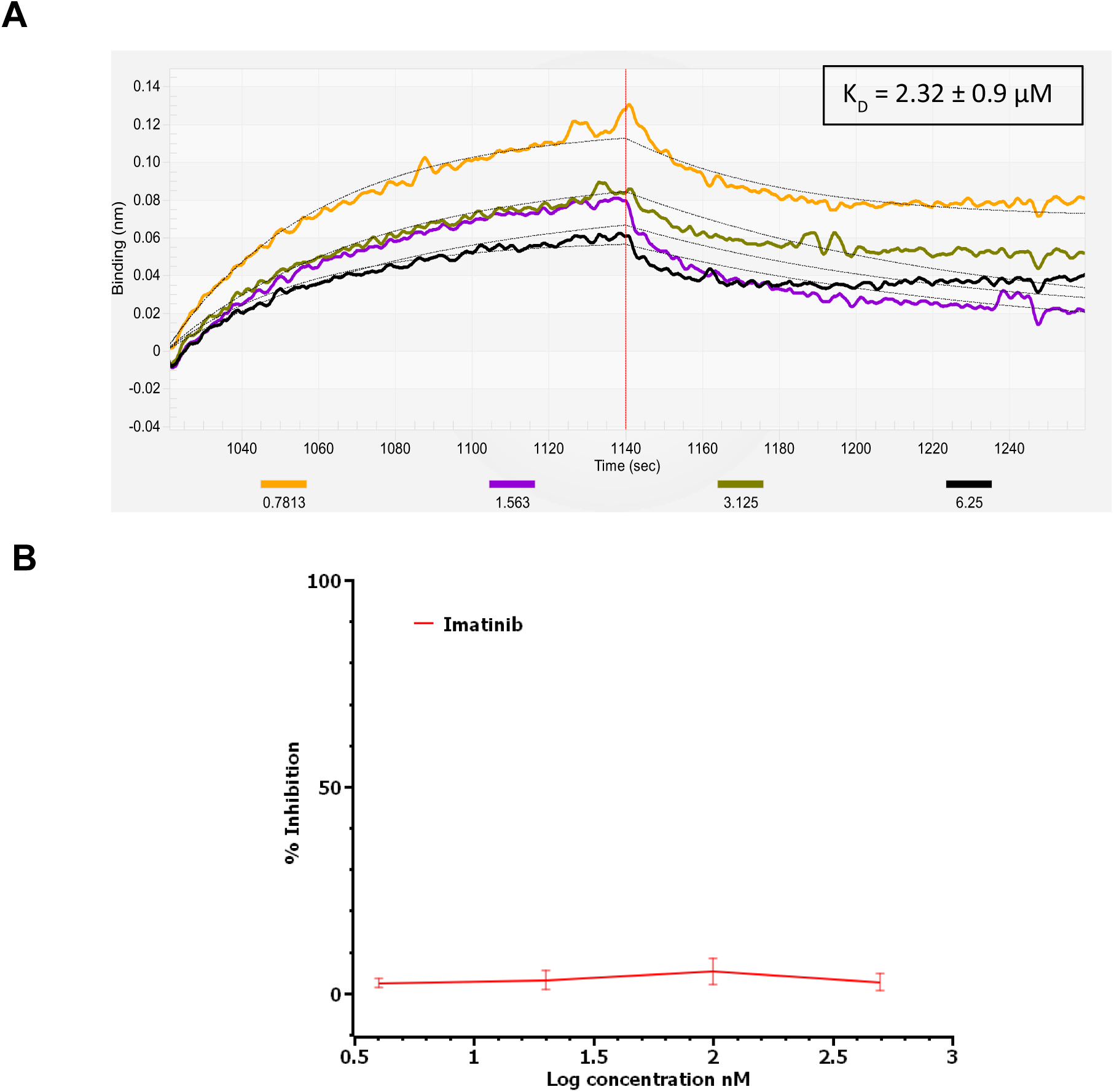
**A]** The association and dissociation curves obtained by BLI reflecting the binding of imatinib to immobilized SARS-CoV-2 RBD protein. Data fitted using the 1:1 binding model are shown in black. **B]** *In vitro* colorimetric assay for evaluation of spike RBD (SARS-CoV-2):ACE2 interaction inhibition in the presence of imatinib. Here, 5-fold dilution of imatinib were done in triplicate. SARS-CoV-2 spike RBD (1 μg/ml) and ACE2 (1 μg/ml) were incubated for 1 hour at room temperature with slow shaking in the presence of various imatinib concentrations. Streptavidin HRP (1:1,000) was added to the reaction mixture. Colorimetric substrate was added to initiate the chromogenic reaction, and 2 minutes were allowed for color development. The reaction was terminated with the addition of 1N HCl and absorbance was measured at 450 nm. Positive control (no inhibitor) was assumed to represent 0% inhibition. Values obtained from test wells (with imatinib) compared to the positive control showed 0% inhibition of RBD:ACE2 interaction, indicating that imatinib does not inhibit spike fusion by direct inhibition.

### Imatinib does not directly inhibit SARS-CoV-2 RBD:ACE2 interaction

*In vitro* colorimetric assays were performed over a 160 pM to 500 nM range in imatinib concentration to assess the ability of imatinib to directly inhibit the RBD:ACE2 interaction. The colorimetric signal of the positive control (no inhibitor) reaction was strong, and the blank wells exhibited an absorbance of ∼ 0.05 at 450 nm as per the manufacturer’s instructions. The test wells (with inhibitor) showed absorbance comparable to the positive control wells indicating that imatinib did not affect the SARS-CoV-2 RBD:ACE2 interaction in the indirect competitive enzyme-linked immunosorbent assay (ELISA), as shown in Fig. 3B.

### Pharmacophore overlap for Abl2 and SARS-CoV-2 spike RBD

A pharmacophore analysis was done to evaluate as to why imatinib showed promising *in-silico* results yet failed to directly inhibit the RBD:ACE2 interaction. The primary binding site of the SARS-CoV-2 RBD was revealed via docking. Pharmacophore analyses were done to further elucidate the interactions between the drug molecules and their receptors (Fig. 4). Twenty-five pharmacophores were collected from the top five binding positions of imatinib at the primary active site of Abl2 tyrosine kinase (native receptor), where each purple sphere represents a pharmacophore as depicted in Fig 4A. Similarly, an additional 25 pharmacophores were collected from the first five binding sites of imatinib at the primary active site of the SARS-CoV-2 RBD. The RBD pharmacophores were represented as yellow spheres in Fig 4B. From the results of ELIXIR-A alignment, it is evident that four pharmacophores between Abl2 kinase and RBD overlapped (red spheres). This significant overlap reveals why the compounds that were originally screened using SARS-CoV-2 RBD also bound to Abl2 kinase, ultimately ensuing in the inhibitory action. The above point is further explicated due to the 54.55% identity between the active sites of the Abl2 (UniProtKB: P42684 aa 288-539) and SARS-CoV-2 spike RBD (UniProtKB: P0DTC2 aa 319-541) generated by a protein BLAST (blastp) [16, 17] search.

**Fig. 4.**
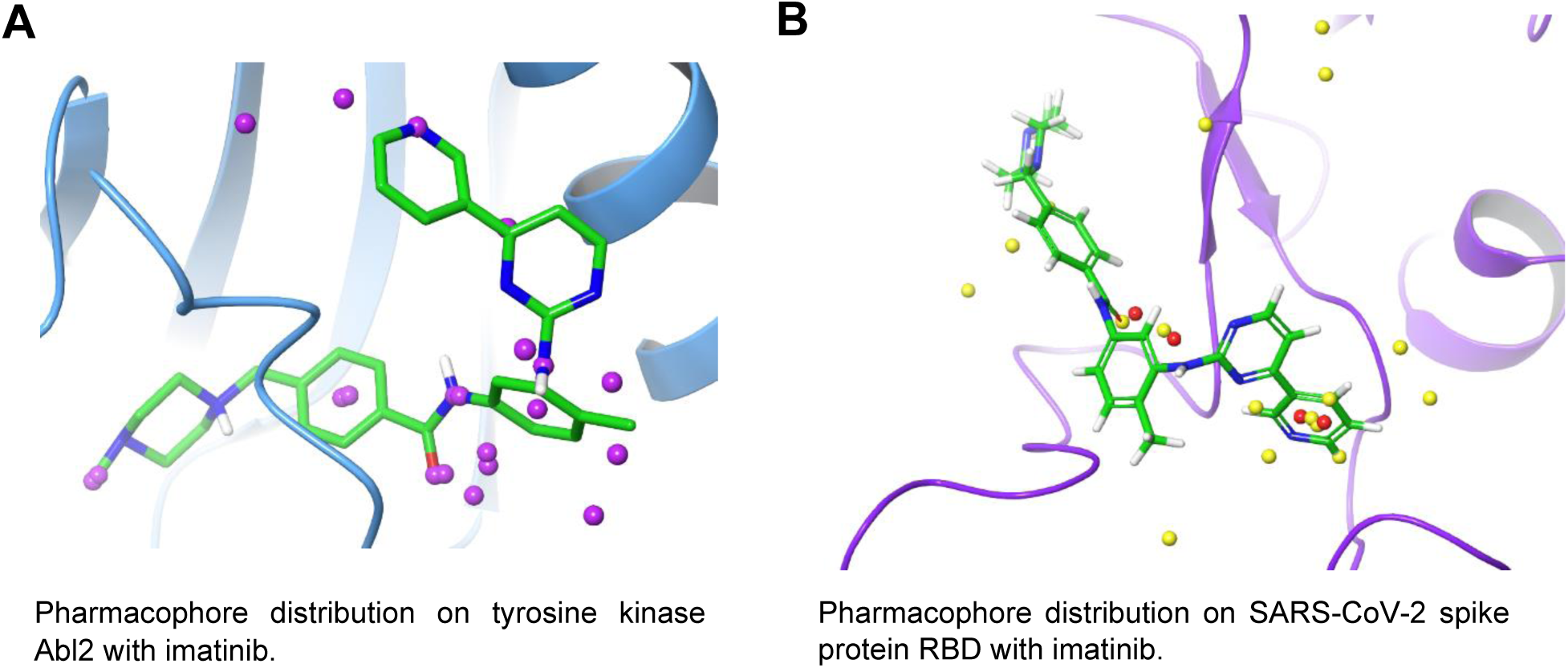
**A]** Pharmacophore distribution of five most stable conformations on tyrosine kinase Abl2 (purple spheres); and **B]** Pharmacophore distribution (yellow spheres) on SARS-CoV-2 spike protein RBD with pharmacophores common to both receptors depicted in red.

## Discussion

There is an urgent need for finding a treatment against the current pandemic of the SARS-CoV-2. Health experts across the globe are trying to use existing clinically approved drugs to treat patients until a specific drug is developed. The present study, using a combination of computational techniques followed by *in vitro* studies, identified imatinib, an FDA approved anti-cancer drug as a potential treatment of SARS-CoV-2 infection. The data indicate inhibition of SARS-CoV-2 replication at IC_50_ of 130 nM. Our results suggest that imatinib prevents viral replication by inhibiting the virus at the fusion stage, possibly by preventing endosomal entry. Binding studies revealed that the affinity of imatinib for the SARS-CoV-2 spike RBD protein is still lower (higher K_D_ value) than the previously published values of nanomolar range (Ligand ID: BDBM13530) [18] for imatinib on Abl tyrosine kinase [19] and in range with the micromolar affinities of imatinib to the SRC-family kinases, FRK and FYN [20]. Although imatinib is not a promiscuous drug, it has been found to bind tightly to tyrosine kinases other than Abl [20, 21]. Pharmacophore mapping between Abl2 and SARS-CoV-2 RBD and a 54.55% identity at the active site of the two proteins explains why imatinib binds to the SARS-CoV-2 RBD as well. However, imatinib failed to directly inhibit the SARS-CoV-2 spike RBD:ACE2 interaction in the competitive ELSA assays. Therefore, it is likely that imatinib causes inhibition of virus fusion via cellular kinase pathway resulting in inhibition of virus replication, as previously described for other coronaviruses [12]. The results provide further evidence supporting the recent clinical trials (ClinicalTrials.gov Identifier: NCT04346147, NCT04357613, NCT04356495, NCT04394416, and NCT04422678) for COVID-19 patients with imatinib.

## Materials and Methods

### Protein Structure Prediction and Validation

A SWISS-MODEL server [22] was used to construct a homology model of the SARS-CoV-2 spike protein using the crystal structure of the SARS-CoV-2 spike protein (PDB:6VSB_chain A) as the template [10]. The genome sequence Wuhan-Hu-1 (GenBank: MN908947.3) was used as a representative of the SARS-CoV-2. Spike protein sequence (GenBank: QHD43416.1) was used as the target sequence [23]. The SWISS-MODEL Structure Assessment Tool was used to validate the quality of the predicted model.

### Molecular Docking

Around 5,800 compounds, including 3,700 nucleoside-like compounds from the Enamine Targeted Antiviral Library (enamine.net) and 2,100 Food and Drug Administration (FDA)-approved drugs from the ZINC15 database [24] were used for molecular docking. All molecules were prepared with obabel [25] from .sdf or .mol2 format to .pdbqt format. The 3D compound structures from the Enamine library were resolved by obabel --gen3d command. The docking file of the protein model was prepared with MGLTools v1.5.4 [26] and the molecules were docked at the RBD of the spike protein via Autodock Vina 1.1.2 [27]. The grid box of 40×60×30 size with 1.0 Å spacing was fixed around the RBD (Thr323-Val511) of the spike protein. Each docking was done in three replicates, and the conformation with the highest binding score was recorded. The batch processing of docking and data collection was performed using an in-house python script which is deposited in GitHub. Data were analyzed statistically using R studio [28] and graphs were constructed with ggplot in R [29]. The ligand-receptor interactions were studied using Schrödinger Maestro [30], and molecules with high docking scores were selected from each screening library for further studies.

### Expression plasmids and cloning

Codon-optimized MERS-CoV (isolate EMC, VG40069-G-N) and SARS-CoV (isolate CUHK-W1; VG40150-G-N) S expression plasmids (pCMV) were ordered from Sino-Biological and subcloned into pCAGGS using the ClaI and KpnI sites. The last 19 amino acids of the SARS-CoV spike protein were deleted to enhance pseudovirus production. Codon-optimized cDNA encoding SARS-CoV-2 S glycoprotein (isolate Wuhan-Hu-1) with a C-terminal 19 amino acid deletion was synthesized and cloned into pCAGSS in between the EcoRI and BglII sites. pVSV-eGFP-dG (#31842), pMD2.G (#12259), pCAG-VSV-P (#64088), pCAG-VSV-L (#64085), pCAG-VSV-N (#64087) and pCAGGS-T7Opt (#65974) were ordered from Addgene. S expressing pCAGGS vectors were used for the production of pseudoviruses, as described below. The cDNA encoding human TMPRSS2 (NM_005656; OHu13675D) was obtained from Genscript. The cDNA fused to a C-terminal HA tag was subcloned into pQXCIH (Clontech) in between the NotI and PacI sites to obtain the pQXCIH-TMPRRS2-HA vector.

### Vero-TMPRSS2 cell line production

Vero-TMPRSS2 cells were produced by retroviral transduction. To produce the retrovirus, 10 μg pQXCIH-TMPRRS2-HA was co-transfected with polyethylenimine (PEI) with 6.5 μg pBS-gag-pol (Addgene #35614) and 5 μg pMD2.G in a 10 cm dish of 70% confluent HEK-293T cells in Opti-MEM I (1X) + GlutaMAX. Retroviral particles were harvested at 72 hours post-transfection, cleared by centrifugation at 2000 x g, filtered through a 0.45μm low protein-binding filter (Millipore), and used to transduce Vero cells. Polybrene (Sigma) was added at a concentration of 4 μg/ml to enhance transduction efficiency. Transduced cells were selected with hygromycin B (Invitrogen).

### Cell lines

HEK-293T cells were maintained in Dulbecco’s Modified Eagle’s Medium (DMEM, Gibco) supplemented with 10% fetal bovine serum (FBS), 1X non-essential amino acids (Lonza), 1mM sodium pyruvate (Gibco), 2mM L-glutamine (Lonza), 100 μg/ml streptomycin (Lonza) and 100 U/ml penicillin. Vero, Vero-TMPRSS2, and VeroE6 cells were maintained in DMEM supplemented with 10% FBS, 1.5 mg/ml sodium bicarbonate (Lonza), 10mM HEPES (Lonza), 2mM L-glutamine, 100 μg/ml streptomycin and 100 U/ml penicillin. All cell lines were maintained at 37°C in a 5% CO_2_, humidified incubator.

### VSV delta G rescue

The protocol for VSV-G pseudovirus rescue was adapted from Whelan and colleagues (1995). Briefly, a 70% confluent 10 cm dish of HEK-293T cells was transfected with 10μg pVSV-eGFP-dG, 2μg pCAG-VSV-N (nucleocapsid), 2μg pCAG-VSV-L (polymerase), 2μg pMD2.G (glycoprotein, VSV-G), 2μg pCAG-VSV-P (phosphoprotein) and 2μg pCAGGS-T7Opt (T7 RNA polymerase) using PEI at a ratio of 1:3 (DNA:PEI) in Opti-MEM I (1X) + GlutaMAX. Forty-eight hours post-transfection the supernatant was transferred onto new plates transfected 24 hours prior with VSV-G. After a further 48 hours, these plates were retransfected with VSV-G. After 24 hours the resulting pseudoviruses were collected, cleared by centrifugation at 2000 x g for 5 minutes, and stored at −80°C. Subsequent VSV-G pseudovirus batches were produced by infecting VSV-G transfected HEK-293T cells with VSV-G pseudovirus at a MOI of 0.1. Titres were determined by preparing 10-fold serial dilutions in Opti-MEM I (1X) + GlutaMAX. Aliquots of each dilution were added to monolayers of 2 × 10^4^ Vero cells in the same medium in a 96-well plate. Three replicates were performed per pseudovirus stock. Plates were incubated at 37°C overnight and then scanned using an Amersham Typhoon scanner (GE Healthcare). Individual infected cells were quantified using ImageQuant TL software (GE Healthcare). All pseudovirus work was performed in a Class II Biosafety Cabinet under BSL-2 conditions at Erasmus Medical Center.

### Coronavirus S pseudovirus production

For the production of MERS-CoV, SARS-CoV, and SARS-CoV-2 S pseudovirus, HEK-293T cells were transfected with 10 μg S expression plasmids. Twenty-four hours post-transfection, the medium was replaced for in Opti-MEM I (1X) + GlutaMAX, and cells were infected at a MOI of 1 with VSV-G pseudotyped virus. Two hours post-infection, cells were washed three times with OptiMEM and replaced with medium containing anti-VSV-G neutralizing antibody (clone 8G5F11; Absolute Antibody) at a dilution of 1:50,000 to block remaining VSV-G pseudovirus. The supernatant was collected after 24 hours, cleared by centrifugation at 2000 x g for 5 minutes and stored at 4°C until use within 7 days. Coronavirus S pseudovirus was titrated on VeroE6 cells as described above.

### Pseudovirus assay

Transduction experiments were carried out by incubating pseudovirus with imatinib at concentrations ranging from 0-125nM in Opti-MEM I (1X) + GlutaMAX for 1 hour at 37°C. Pseudovirus-imatinib mixes were added to monolayers of 2 × 10^4^ Vero or Vero-TMPRSS2 cells in a 96-well plate. Plates were incubated for 16 hours before quantifying GFP-positive cells using an Amersham Typhoon scanner and ImageQuant TL software.

### *In vitro* toxicity of imatinib

To determine the toxicity profile of imatinib, we performed the MTT assay using a 1-hr and an 8-hr design. Briefly, a serial dilution of imatinib was prepared and incubated on Vero cells for 1 hr at 37°C. Subsequently, cells were washed, further cultured for eight hrs. In the 8-hr design, cells were incubated with a serial dilution of imatinib for eight hours without a washing step.

### *In vitro* efficacy of imatinib

We tested serial dilutions of imatinib for its ability to neutralize SARS-CoV-2 (German isolate; GISAID ID EPI_ISL 406862; European Virus Archive Global #026V-03883) using a plaque reduction neutralization test (PRNT) as previously described [31]. Fifty μL of the virus suspension (200 spot forming units) was added to each well and incubated at 37°C for either 1 hr. Following incubation, the mixtures were added on Vero cells and incubated at 37°C for either 1hr or 8 hrs. The cells incubated for 1 hr were then washed and further incubated in medium for 8 hrs. After the incubation, the cells were fixed and stained with a polyclonal rabbit anti-SARS-CoV antibody (Sino Biological; 1:500). Staining was developed using a rabbit anti-SARS-CoV serum and a secondary alexa-fluor-labeled conjugate (Dako). The number of infected cells per well were counted using the ImageQuant TL software.

### BLI

The binding kinetics of imatinib on SARS-CoV-2 RBD protein were studied using a BLItz® system (FortéBio). Experiments were conducted using the advanced kinetics mode, at room temperature and a buffer system consisting of 1X Kinetics Buffer (FortéBio), 5% anhydrous dimethyl sulfoxide (DMSO; Sigma Aldrich). Recombinant His-tagged SARS-CoV-2 RBD protein (40592-V08H; Sino Biological) at a concentration of 10 μg/ml was loaded on Anti-Penta-HIS (HIS1K) Biosensors (FortéBio), followed by a washing step with assay buffer to block the unoccupied sensor surface. The association and dissociation profiles of imatinib (Sigma Aldrich) were measured at various concentrations (four-point serial dilutions from 6.25 μM to 0.78 μM). A reference biosensor loaded in the same manner with 0 μM imatinib was used for baseline correction in each assay. The final binding curves were analyzed with the BLItz Pro 1.3 Software (FortéBio) using the 1:1 global-fitting model. The assay was repeated twice to validate the binding constants. Here, data is represented as mean ± SD.

Similarly, SARS-CoV-2 RBD was immobilized on HIS1K biosensors to study the binding kinetics of mFc-tagged hACE2 (10108-H05H; Sino Biological) before being dipped into tubes containing the 1X Kinetics buffer. Various concentrations of ACE2 (four-point serial dilutions from 800 to 100 nM) were used to measure the association and dissociation profiles. Data were reference subtracted and fit to a 1:1 binding model using the BLItz Pro 1.3 Software.

### *In vitro* colorimetric assay for RBD (SARS-CoV-2):ACE2 interaction inhibition

The ability of imatinib to inhibit the interaction of spike RBD:ACE2 proteins was evaluated by using the Spike RBD (SARS-CoV-2): ACE2 Inhibitor Screening Colorimetric Assay Kit (BPS Bioscience). The Avi-His-tagged Spike S1 RBD (SARS-CoV-2) protein 50 ng/well in PBS was coated onto 96-well microplate by overnight incubation at 4°C. Blocking Buffer 2 was used to block the nonspecific binding sites by incubation for 1 hour. Different concentrations of imatinib were added and incubated for 1 hour at room temperature with slow shaking. For the wells designated “Blank” and “Positive Control”, inhibitor buffer (PBS with 0.5% DMSO) was added. The reaction was initiated by adding ACE2 His-Avi-tagged Biotin-labeled HiP™ protein (50 ng/well) in 1X Immuno Buffer 1 to the “Positive Control” and “Test Inhibitor” wells by incubation for 1 hour at room temperature with slow shaking. Streptavidin-HRP (dilution 1:1,000 in Blocking Buffer 2) was added to each well and incubated at room temperature for 1 hour with slow shaking. Washing procedure (3 × 100 μl 1X Immuno Buffer 1) was performed after each step. The chromogenic reaction was initiated by adding Colorimetric HRP substrate to each well and incubated at room temperature until blue color was developed (approximately 2 minutes) in the “Positive Control” well. After the blue color was developed, the reaction was terminated by adding 1N HCl, and absorbance at 450 nm was measured using Synergy H1 Hybrid Multi-Mode Microplate Reader (BioTek Instruments).

### Pharmacophore overlay analysis

The pharmacophore model was generated using Pharmit [32], an online interactive platform to elucidate pharmacophores from the receptor and ligand complex. Top five binding conformations of drug-protein complexes were produced by Autodock Vina. The pharmacophores of the ligands interacting with the receptor were considered active pharmacophores while the rest were defined as inactive pharmacophores. The pharmacophores from the native receptor of imatinib (Abl2 tyrosine kinase; PDB:3GVU) and RBD of SARS CoV-2 were generated using imatinib as the ligand. Using the Enhanced Ligand Exploration and Interaction Recognition Algorithm (ELIXIR-A), the two sets of pharmacophores were merged and processed to identify any overlap in 3D space. A detailed description of ELIXIR-A can be found in our previous work [33] and the algorithm has been deposited in GitHub.

## Code availability

The python script ‘ELIXIR-A-Vina-Batch-Screening-Module’ used for running docking jobs in batch mode and ELIXIR-A, the algorithm used for pharmacophore mapping have been deposited in GitHub and will be made public upon publication of the manuscript.

## Acknowledgment

We gratefully acknowledge the support from Texas A&M High Performance Research Computing (HPRC) and TAMU Laboratory for Molecular Simulation (LMS). We would like to thank Dr. Lisa Perez (Associate Director for Advanced Computing Enablement, HPRC TAMU) for guidance with MD simulations. We are thankful to Mart Lammers for allowing us to use the fusion assays

## Funding

DR was supported by the Ministry of Health of the Czech Republic (project No. 20-05-00472).

## Author contributions

SF, BM, and DR conceived and designed the study. First author NM designed the experiments, performed BLI studies and immunoassays, reviewed literature, and compiled the manuscript and figures. Co-first author HW conducted *in silico* experiments and compiled figures. SM did literature review on the resulting compounds and compiled the manuscript and figures. BM performed virology experiments. SF directed and verified studies and authored the manuscript. All authors reviewed and edited the paper.

## Competing interests

All the authors declare that there are no conflicts of interest.

## Notes

### Competing Interest Statement

The authors have declared no competing interest.

