## Supplementary Data for "Bcr-Abl tyrosine kinase inhibitor imatinib as a potential drug for COVID-19"

**Affiliations**
[1]Biological and Agricultural Engineering Department, Texas A&M University, College Station, TX 77843, USA.
[2]Artemis One Health Research Institute, Delft, The Netherlands.
[3]Veterinary Research Institute, Brno, and Institute of Parasitology, Biology Centre of the Czech Academy of Sciences, Ceske Budejovice, Czech Republic.

[#]These authors contributed equally.

[*]**Corresponding authors**
Dr. Sandun Fernando:
Dr. Byron Martina:

**This PDF file includes:**

Tables S1 and S2
Figure. S1
Caption for Movie S1
References

**Table S1. Vina docking scores of the selected compounds.**

| **Compounds** | **SARS-CoV template (PDB: 6ACD_chain A)** | **SARS-CoV-2 template (PDB: 6VSB_chain A)** |
|---|---|---|
| | **Docking Score (kcal/mol)** | |
| DMSO | -1.90 ± 0.00 | -2.10 ± 0.00 |
| Antiviral825 | -6.90 ± 0.00 | -6.30 ± 0.00 |
| Antiviral2038 | -6.30 ± 0.10 | -6.20 ± 0.20 |
| Antiviral2981 | -6.80 ± 0.00 | -6.00 ± 0.00 |
| ZINCFDA130 (ergotamine) | -8.00 ± 0.00 | -7.70 ± 0.00 |
| ZINCFDA515 (ponatinib) | -8.47 ± 0.06 | -7.63 ± 0.06 |
| ZINCFDA754 (imatinib) | -7.50 ± 0.10 | -6.80 ± 0.17 |
| ZINCFDA2083(glecaprevir) | -7.10 ± 0.17 | -7.20 ± 0.35 |

Note:Docking was done in triplicate for each molecule and data are represented as mean ± standard deviation.

**Table S2 – Information of substances screened from initial docking studies.**

| **Substance** | **2D structure** | **Applications and mechanism of action** | **References** |
|---|---|---|---|
| ZINCFDA754/ imatinib/ ZINC000019632618 | 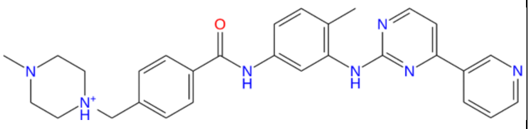  | Leukemia treatment by inhibition of Bcr-Abl tyrosine kinase. | [1] |
| Antiviral2038 / Z787722876 / ZINC50038784 | 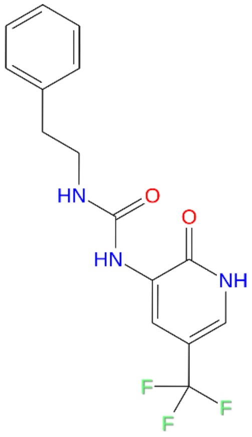  | No reported bioactivity. Antiviral properties based on Enamine predictions. | |

| | | | |
|---|---|---|---|
| Antiviral2981 / Z1452532074 / ZINC170674881 | 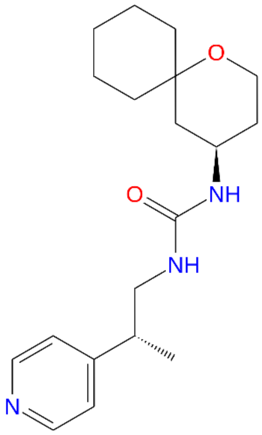 | Has a molinspiration bioactivity score of 0.35 as GPCR ligand | [1] |
| Antiviral825 / Z1277226201 / ZINC104169890 | 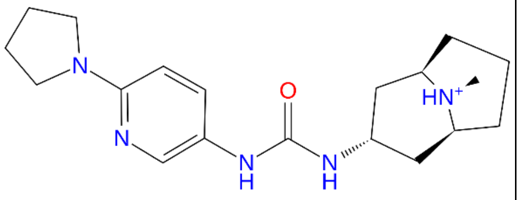 | Has good molinspiration bioactivity scores as GPCR ligand (0.43), Ion channel modulator (0.38), kinase inhibitor (0.27) and enzyme inhibitor (0.2) | [1] |
| ZINCFDA130 / ergotamine/ ZINC000052955754 | 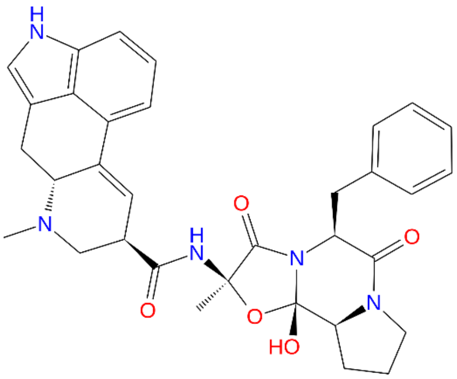 | Migraine treatment via acting as an agonist to 5-HT1A, 5-HT1B, 5-HT1D, and 5-HT1F receptors | [2] |
| ZINCFDA2083 / glecaprevir/ ZINC164528615 | 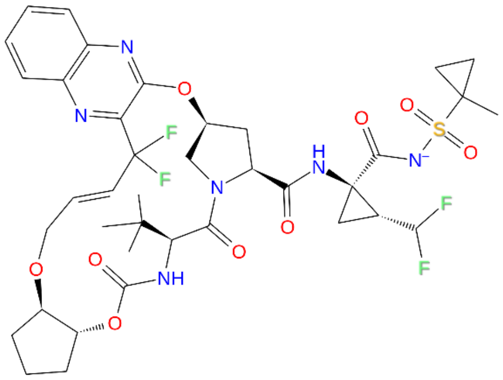 | Hepatitis C treatment by NS3/4 protease inhibition | [3] |

| | | | |
|---|---|---|---|
| ZINCFDA515 / ponatinib/ ZINC000036701290 | 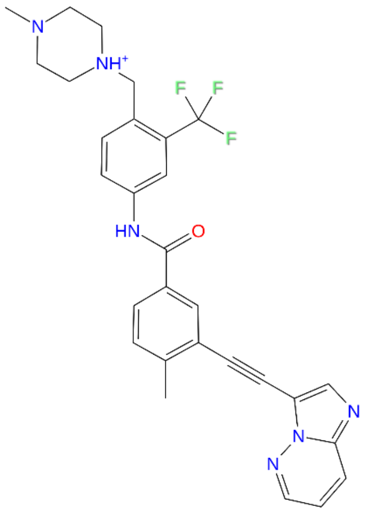 | Leukemia treatment by inhibition of Bcr-Abl tyrosine kinase | [4] |

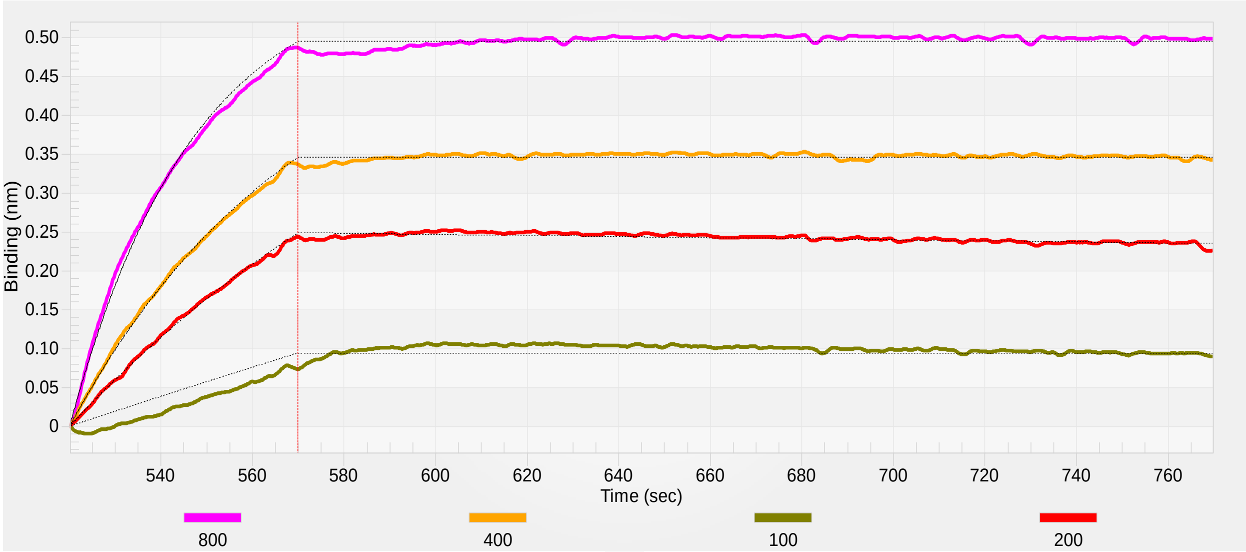


**Fig. S1** Binding kinetics of ACE2 (mFc tag) at concentrations 100-800 nM onto immobilized SARS-CoV-2 RBD (his tag) via HIS1K biosensors. $K_D$ = 5.936 nM, $k_a = 4.768 \times 10^4$ $M^{-1}s^{-1}$, $k_d = 2.83 \times 10^{-4}$ $s^{-1}$. Data fitted using the 1:1 binding model are shown in black.

**Movie S1.** Molecular dynamics simulation video of SARS-CoV-2 spike protein (RBD) interacting with imatinib.
